# LARGE MICROBIOTA SURVEY REVEALS HOW THE MICROBIAL ECOLOGY OF COOKED HAM IS SHAPED BY DIFFERENT PROCESSING STEPS

**DOI:** 10.1101/865857

**Authors:** Marine Zagdoun, Gwendoline Coeuret, Méry N’Dione, Marie-Christine Champomier-Vergès, Stéphane Chaillou

## Abstract

The production of cooked ham involves numerous steps shaping the microbial communities of the final product, with consequences on spoilage metabolites production. To identify the main factors driving the ecology of ham and its spoilage, we designed a study encompassing five variables related to ham production: type of storage during meat transportation, churning speed, drain-off time, slicing line and O_2_ packaging permeability. We obtained about 200 samples from the same facility and we characterized *i)* their microbiota based on *gyrB* amplicon sequencing *ii*) their production of spoilage-related metabolites based on E-Nose analysis and enzymatic assays. The slicing was the most critical step, shaping two general types of microbiota according to the slicing line: one dominated by *Carnobacterium divergens* and another one dominated by *Leuconostoc carnosum* and *Serratia proteamaculans*. Regarding metabolites production, *L. carnosum* was associated to D-lactic acid, ethanol and acetic acid production, whereas *Serratia proteamaculans* was associated to acetic acid production. This last species prevailed with highly O_2_-permeable packaging. Within a given slicing line, we observed campaign-based variations, with *Lactobacillus sakei*, *Leuconostoc mesenteroides* and *Carnobacterium maltaromaticum* prevalent in summer. *L. sakei* was associated with L-lactic acid production and *C. maltaromaticum* with formic and acetic acid productions.

## 1. INTRODUCTION

The production of cooked ham involves many processing steps, with each being a potential source of bacterial contamination or re-contamination post-cooking. As a consequence, the shelf-life of this meat product is strongly dependent on the microbiota dynamic through these various steps. It is also related with how the process parameters will influence the metabolic activities of the growing bacterial species during storage (Audenaert et al., 2010; Dušková et al., 2016; Han et al., 2011; Raimondi et al., 2019; Piotrowska-Cyplik et al., 2017; Vasilopoulos et al., 2010).

The variability of ham begins at the most upstream step of the process as initial raw pork cuts usually contain diverse microbiota depending on the slaughtering and butchering process (De Filippis et al., 2013; Wheatley et al., 2014). Bacterial contamination usually increases in the processing plants through the contamination of surfaces and tools (Stellato et al., 2016) along the various processing steps (Hultman et al., 2015; Samelis et al., 1998), as well as with ingredients used in the brine such as spices (Benson et al., 2014). Furthermore, the efficiency of brining requires a churning incubation time, a step which was shown to participate in the selection of some bacterial species, such as *Brochothrix thermosphacta* or *Carnobacterium divergens* (Samelis et al., 1998). After churning, bacterial load is close to 10^5^ cfu/g (Dušková et al., 2016; Samelis et al., 1998; Vasilopoulos et al., 2010). The following cooking step at 70°C generally reduces the bacterial load back to practically zero (Dušková et al., 2016; Vasilopoulos et al., 2010). However, some studies showed that thermoduric strains such as *Leuconostoc carnosum* or *Weissella viridescens* could resist cooking (Björkroth et al., 1998; Comi and Iacumin, 2012). Other studies showed that *Leuconostoc mesenteroides* and *L. carnosum* were also brought by the post-cooking downstream processing steps, either through the hands of worker while de-molding, via the slicing machine, the packaging or even via the air (Björkroth and Korkeala, 1997; Dušková et al., 2016; Mol et al., 1971; Samelis et al., 1998). Then, depending on the storage temperatures and the type of packaging used, a final reduction of bacterial diversity takes place, selecting psychrotrophic anaerobic microbiota (Audenaert et al., 2010; Chaillou et al., 2015), the activity of which will eventually cause spoilage.

Thus, if the microbiota of ham after cooking does probably result of both surviving microbiota after cooking and post-cooking contaminations, it is yet unclear to what extent raw meat and pre-cooking process steps can shape the final microbiota of ham. In this line of view, Geeraerts and co-workers (2017) have nevertheless underlined the variability of bacterial communities of hams between different facilities. They have also shown that those communities can vary according to the sampling period. Even hams produced within a same facility revealed to harbor variable communities (Audenaert et al., 2010; Vasilopoulos et al., 2010, 2008). More recently, Raimondi and co-workers (2019) described a wide variability in the microbial assemblages of cooked ham from ten different producers in Europe. Despite the low number of ham analyzed in this study (14 different ham in total), their work showed that the quality of hams at the end of the storage period, as measured by organic volatile compounds analysis, was highly dependent on the microbiota present on these products.

Given this complex processing, the spoilage microbial ecology of ham remains difficult to decipher and it is hard for the meat industry to implement useful support decision tool for process management and prevention of food losses. To us, the works described above represent a revealing illustration of the necessity of adopting a large sampling strategy and robust statistical analysis in order to capture the overwhelming variability created by the many steps of cooked ham processing. The aim of this study was to provide such type of large-scale approach based on a factorial design to allow the analysis of variations within five main processing steps. We focused our analysis on a single processing factory, located in France, and by designing a large sampling (over 200 samples), spanning a production period of six months. To improve our accuracy in tracing bacterial taxa throughout processing, we performed *gyrB* amplicon sequencing, a powerful alternative to 16S rRNA gene sequencing for accurate bacterial identification at the intra-species level (Poirier et al., 2018b). We then combined this diversity analysis with the quantification of pH and of five metabolites (ethanol, D- and L-lactate, acetate, formate), used as indicators of fermentation end-products contributing to the pattern of ham spoilage.

## 2. MATERIALS AND METHODS

### 2.1. Industrial ham production

Raw pork meat for the production of ham was obtained from a single slaughterhouse. After deboning, raw meat was transported to the ham production factory (see processing step in Table 1). Meat was then brined to reach a final concentration (wt/wt) in the product of 1.4% sodium chloride, 0.5% glucose, 0.08% sodium nitrite, and 0.03% sodium ascorbate. Each sub-batch (determined by variations in processing parameters; see Table 1) was then churned and molded into individual loaves and cooked until the center reached 70°C. After cooking, draining-off, and slicing, hams were packed under 50% CO_2_/50% N_2_ modified-atmosphere conditions.

**TABLE 1.** Levels of the different processing steps

| Processing step | Description of the step variations | Labeling of levels | Description of the levels analyzed in this study |
| --- | --- | --- | --- |
| Meat transportation | Time and storage conditions of the transportation of pork meat from slaughterhouse to the ham factory may vary as a function of meat origin. | Vacuum | Storage in vacuum packaging for ten days at 4°C |
|  |  | Air | Storage in open air for three days at 4°C |
| Churning | During ham production, the number of churning rotations is a constant but pause time between two rotation sessions may vary in response to the arrival of meat batches and staff organization. | Quick | Standard quick churning at 4°C, carried out during the workweek |
|  |  | Slow | Slow churning (four times longer than standard churning) at 4°C, carried out during the weekend |
| Drain-off | The drain-off time after cooking may be modified in response to production demand. | Long | Drain-off for twelve days at 4°C when production demand is low. |
|  |  | Short | Drain-off for four days at 4°C when production demand is high. |
| Slicing line | Several types of slicing lines may be present within a production unit. | Line A | First-generation slicing technology. |
|  |  | Line B | Second-generation slicing technology (more complex equipment). |
| O<sub>2</sub> packaging permeability | The film used for sealing the modified atmosphere packaging may vary in its degree of permeability to O<sub>2</sub>. | High O<sub>2</sub> | Single-layer APET packaging (amorphous polyethylene terephthalate) has an O<sub>2</sub> permeability of 16.3 cm<sup>3</sup>.m<sup>-2</sup>.day.10<sup>5</sup> Pa. |
|  |  | Low O<sub>2</sub> | Multiple-layer polyvinyl chloride/vinyl alcohol/polyethylene packaging has an O<sub>2</sub> permeability of 1.24 cm<sup>3</sup>.m<sup>-2</sup>.day.10<sup>5</sup> Pa. |

### 2.2. Ham storage conditions

Industrial hams were sent to our laboratory immediately after packaging using refrigerated transportation (2 days at 4°C). Hams were stored in FRIOCELL 55 refrigerated incubators (MMM Medcenter, Planegg, Germany) at 4°C for 5 more days, then at 8°C until the Use-By-Date (UBD) or for 15 more days (past UBD). Control samples (T_0_) were immediately frozen at −20°C after packaging and sent frozen to our laboratory. At the end of storage, all ham samples were frozen at −20°C until pH measurement, DNA extraction, and recovery of the microbiota were performed.

### 2.3. Gas and pH measurements

O_2_ and CO_2_ levels were quantified using an Oxybaby probe (WITTGAS, Morsang-sur-Orge, France). Then, pH was measured using a FiveEasy FE20 surface pH probe (Mettler Toledo AG, Schwerzenbach, Schwitzerland). The pH data correspond to a mean of five measurements per sample at various areas on the slices.

### 2.4. Identification and relative quantification of volatile organic compounds

Volatile compounds in ham samples were detected with a Heracles II electronic nose (Alpha MOS, Toulouse, France), using ultra-fast gas chromatography. Samples were analyzed on two independent columns of different polarity (nonpolar MXT-5 and medium polar MXT-1701, length 10 m each) for an accurate identification of compounds using their KOVAT number. Agrochembase, an internal database of AlphaMOS, provided information on chemical analysis and sensorial olfactory descriptors.

### 2.5. Microbiota recovery and bacterial load quantification

For each ham sample, four independent DNA extractions were performed (each of one quarter of a ham) and were subsequently pooled. Our protocol was based on that of Fougy and colleagues (Fougy et al., 2016), adapted as follows: 20 g of meat were homogenized for 2 × 30 s in a stomacher bag (BagPage, Interscience, Saint-Nom-la-Bretèche, France) with 40 mL of sterile ultrapure water supplemented with 1% Tween 80 (Acros Organics, Waltham, USA). Large meat debris were filtered from the bag and 33 mL of the microbe-containing ham solution was collected. One milliliter was reserved to estimate viable bacterial counts, and the remaining 32 mL were centrifuged at 500 × *g* for 3 min at 4°C (Allegra X-15R, Beckman Coulter, Villepinte, France) to spin down the remaining meat fibers and debris. A maximum volume of 25 mL of supernatant was then centrifuged at 3000 × *g* for 5 min at 4°C to spin down microbial cells. The supernatant was frozen at −80°C for further metabolite quantification, and the bacterial pellets were washed in 1 mL of sterile ultrapure water, centrifuged at 3000 × *g* for 5 min at 4°C (Centrifuge 5424, Eppendorf, Hamburg, Germany), and frozen at −20°C for bacterial DNA extraction. For quantification of the bacterial population, the reserved 1 mL of ham solution was subjected to decimal dilutions and plated on Columbia agar with 5% sheep blood (Thermo Fisher Scientific, Illkirch Graffenstaden, France) in order to estimate the overall cultivable bacterial load. Bacterial colonies were counted after 72 hours of incubation at 20°C.

### 2.6. Quantification of end-products

Supernatants of ham samples were clarified to eliminate proteins using the Carrez clarification kit (Merck KGaA, Darmstadt, Germany). Proteins were precipitated via the successive addition of 600 µL of Carrez reagent I and 600 µL of Carrez reagent II to 5 mL of supernatant. After adjusting the pH to be between 7.5 and 8.5 (required for enzymatic assays), the samples were centrifuged at 5000 × *g* for 20 min to spin down precipitated proteins. Residual proteins were then eliminated by filtration on 0.22-µm high-protein-retention Ministart syringe filters (Sartorius Stedim Biotech GmbH, Goettingen, Germany). Megazyme enzymatic assays (Megazyme Wicklow, Ireland) were used to quantify concentrations of ethanol (K-ETOH), L- and D-lactic acid (K-LAC), acetic acid (K-ACET), and formic acid (K-FORM), according to the manufacturer’s instructions. For all kits, NADH absorbance was measured using a microplate reader (SYNERGY Mx, BIOTEK, Colmar, France).

All calculations of end-product concentrations took into account the dilution factor (20 g of meat in 40 mL of sterile water with 1% Tween 80 for microbiota recovery, and addition of 1.2 mL of Carrez reagents) in order to estimate the real concentrations in meat.

### 2.7. Bacterial DNA extraction and quality control

Bacterial DNA was extracted with a High Pure PCR Template Preparation kit (Roche Diagnostics Ltd, Burgess Hill, West Sussex, UK), with the following adaptations: for cellular lysis, bacterial pellets were resuspended in 200 µL PBS that contained lysozyme (final concentration of 20 mg.mL^−1^) and Mutanolysin (12U). This solution was incubated at 37°C for 60 min in a water bath (Polystat 86602, Bioblock Scientific, Illkirch, France). Then, 200 µL of binding buffer and 40 µL of Proteinase K were added; the solution was vortexed and incubated at 70°C for 30 min in a second water bath (APELEX, JULABO, Seelbach, Germany). DNA was precipitated and purified on columns according to the manufacturer’s instructions, with the exception of a reduction of the final elution volume to 100 µL in order to concentrate the DNA. Technical replicates for each ham sample were pooled following DNA extraction and frozen at −20°C. After pooling, double-strand DNA quantification was performed with the Qubit dsDNA HS assay kit (Invitrogen, Carlsbad, USA).

### 2.8. PCR protocols for *gyrB* and 16S V3-V4 rRNA gene amplification, purification and quality control of amplicons

The internal fragment of the g*yrB* gene (around 280 bp) was amplified as described in (Barret et al., 2015; Poirier et al., 2018b) with AccuPrime Taq DNA polymerase (Invitrogen, Carlsbad, USA) and F64 and R353 degenerated primers (5’-MGNCCNGSNATGTAYATHGG-3’ and 5’-CNCCRTGNARDCCDCCNGA-3’, respectively). The forward and reverse primers carried the Illumina adapters 5’-CTTTCCCTACACGACGCTCTTCCGATCT-3’ and 5’-GGAGTTCAGACGTGTGCTCTTCCGATCT-3’, respectively. To balance the high degeneration rate of the primers, the final primer concentration was increased to 1000 nM. PCR was performed on 10 ng of each pooled DNA sample using a PTC-200 thermal cycler (Peltier, BioRad, Hercules, USA). Amplification started with an initial denaturation step of 2 min at 94°C. This was followed by 35 cycles of denaturation (94°C for 30 s), annealing (55°C for 60 s), and elongation (68°C for 90 s), and ended with a final elongation step of 10 min at 68°C. PCR bias was minimized by pooling three independent PCR replicates per sample.

To assess potential biases in the recovered microbial diversity, controls using 16Sr V3-V4 rRNA gene amplifications (around 450 bp) were performed on fourteen selected samples; these amplifications used MolTaq 16S DNA polymerase (Invitrogen, Carlsbad, USA) and two primers that targeted the V3 and V4 region (Klindworth et al., 2013; Poirier et al., 2018a), 5’-ACGGRAGGCWGCAGT-3’ and 5’-TACCAGGGTATCTAATCCT-3’. The forward and reverse primers carried the Illumina adapters 5’-CTTTCCCTACACGACGCTCTTCCGATCT-3’ and 5’-GGAGTTCAGACGTGTGCTCTTCCGATCT-3’, respectively. The final primer concentration was 200 nM. PCR was performed on 10 ng of the pooled DNA sample with a PTC-200 thermal cycler, with the following cycling parameters: initial denaturation for 1 min at 94°C, followed by 30 cycles of denaturation (94°C for 60 s), annealing (65°C for 60 s), and elongation (72°C for 60 s), and a final elongation step of 10 min at 72°C. PCR bias was minimized by pooling three independent PCR products per sample.

The *gyrB* and 16S rRNA gene amplicons were purified using a QIAquick kit (Qiagen, Hilden, Germany) and their quality and concentration were checked with a DNA1000 chip (Agilent Technologies, Paris, France).

### 2.9. Illumina MiSeq library preparation and sequencing

In order to pool and sequence a large number of libraries simultaneously, samples were coded with individual barcode sequences during library preparation: 6-bp unique indices and the second part of the forward (5’-AATGATACGGCGACCACCGAGATCTACACT-3’) and reverse adapters (5’-CAAGCAGAAGACGGCATACGAGAT-NNNNNN-GTGACT-3’) were added during the Illumina PCR. PCR comprised an initial denaturation step at 94°C for 10 min, followed by 12 amplification cycles of 94°C for 1 min, 65°C for 1 min, and 72°C for 1 min, and a final elongation step at 72°C for 10 min. Amplicons were then purified on Clean PCR magnetic beds (Clean NA, Alphen aan den Rijn, The Nederlands). Their concentration after purification was estimated with a Nanodrop spectrophotometer (Thermo Scientif, Whaltham, USA) and their quality was checked using a Fragment Analyzer (AATI, Santa Clara, USA). Reads from libraries were pooled in equal amounts, in order to obtain a final concentration for sequencing between 5 and 20 nM. Pools were denatured with NaOH, mixed with the PhiX control library v3 (diluted to 7 pM; Illumina, San Diego, USA), and loaded on an Illumina MiSeq cartridge. The MiSeq Reagent Kit v3 (2×300 paired-end reads, 15 Gb output) was used according to the manufacturer’s instructions. The quality of the obtained sequences was checked with FastQ files generated at the end of the run (MiSeq Reporter Software, Illumina, USA) and additional PhiX Control. The corresponding pairs of sequences were then attributed to their respective samples using the individual multiplexing barcodes.

### 2.10. Quality control of raw MiSeq reads and OTU clustering parameters

Samples were sequenced in three runs (295 amplicons of *gyrB* analyzed in runs 1 and 2 and 28 amplicons of 16S rRNA gene analyzed in run 3). Quality controls indicated a quality score of at least 30 for 91% of the reads and a median number of 44,022 ± 33,312 reads per sample.

We used FastQC (https://github.com/s-andrews/FastQC) to assess sequencing quality. Paired-end sequences were then merged using PEAR v0.9.10 (Zhang et al., 2014) with a minimum overlap of 20 nucleotides for both *gyrB* and 16S rRNA gene and a threshold p-value of 0.01. We retained merged sequences with a size of 280 ± 50 bp for *gyrB* and 450 ± 50 bp for 16S rRNA gene. Primer-binding sequences were removed using CUTADAPT v1.12 (Martin, 2011) with a maximum error rate of 0.01; low-quality bases at the ends of the merged sequences (quality threshold of 20, sequences with ambiguous bases N) were eliminated using SICKLE v1.330 (https://github.com/najoshi/sickle).

Quality-filtered merged sequences were then loaded in the FROGS (Find Rapidly OTUs with Galaxy Solutions) pipeline (Escudié et al., 2018) and dereplicated. SWARM clustering (Mahé et al., 2014) was performed using a maximal aggregation distance of three nucleotides for 16S rRNA gene sequences; for *gyrB* sequences, clustering was more stringent, with a maximal aggregation distance of two nucleotides in order to potentially assign OTUs to the subspecies level. Chimeras were removed using the VCHIM feature of the VSEARCH package (https://github.com/torognes/vsearch). Clusters were further filtered to remove spurious low-abundance OTUs arising from sequencing artifacts. As reported by Bokulich and colleagues (2013), these types of OTUs appear in very few samples with very low abundance (<0.005%). Therefore, for the *gyrB* reads we retained OTUs with at least 100 reads in the whole dataset (0.002% of 5.9 million reads in total), and for the 16S rRNA gene used as controls, we retained OTUs with at least 10 reads in the whole dataset, since there were fewer samples than for *gyrB* (0.0008% of 1.3 million reads in total).

These steps resulted in the retention of 238 *gyrB* OTUs and 88 16S rRNA gene OTUs. An otu_table with the respective normalized abundances of the different OTUs in each sample was constructed for both the *gyrB* and 16S rRNA gene datasets and is available as supplementary dataset S2. The total number of reads per sample was normalized based on the median sequencing depth of all samples.

### 2.11. Taxonomic assignment and phylogenetic tree construction

For certain species of Firmicutes, PCR amplification of *gyrB* can also recover the *parE* gene, which encodes the β subunit of DNA topoisomerase IV. In order to determine which of the two genes had been amplified for each species, the OTU sequences were blasted against the *gyrB*/*parE* databases established by Poirier and colleagues (2018b), using the Blastn+ algorithm (Camacho et al., 2009) for taxonomic assignment. Both *gyrB* and *parE* OTUs were retained in our diversity analysis. We considered taxonomic assignments to be valid when 94% of a sequence matched over 200 bp of coverage; valid matches were assigned identifications at least at the species level and in some cases at the subspecies level.

Sequences of 16S rRNA gene were identified using the Ribosomal Database Project (RDP) Classifier (Wang et al., 2007) and the SILVA 128 SSU databases (Quast et al., 2013). Taxonomic assignment of the different OTUs to genera and, when possible, to the species level was carried out based on a threshold of 97% identity. The Ez-taxon-e database was used to identify uncultured species (Kim et al., 2012) that had not been previously identified with SILVA.

For each dataset, phylogenetic trees were created with SeaView software (Gouy et al., 2010) using the Kimura-2-parameter algorithm (Kimura, 1980) as distance estimator. Distance matrices were converted to trees using the BioNJ neighbor-joining method (Gascuel, 1997).

### 2.12. Alpha/beta diversity and statistical analyses

Bacterial diversity analyses were performed using the phyloseq package (v1.24.2) of R (McMurdie and Holmes, 2013). The relative abundance of bacterial species in samples was plotted with the function plot_composition. A dendrogram based on Bray-Curtis dissimilarity index values was plotted with the function as.dendogram, using the ward.D2 method, and visualized with the function fviz_dend in the gglot2 package (v3.0.0) of R. To visualize the results of principal coordinate analyses (PCoA), samples were first ordered according to Bray-Curtis dissimilarity index values using a non-metric dimensional scaling (NMDS) approach, and then plotted with the function s.class within the ade4 package (v1.7-13) of R (Chessel et al., 1987). Bacterial diversity was compared among different groups of samples with permutational ANOVA, specifically using the adonis function within the vegan package (v2.5-2; https://github.com/vegandevs/vegan) of R. Boxplots were drawn with the ggboxplot function from R’s ggpubr package, v0.1.8. In Figure 5, the significance of differences between groups of samples was assessed with ANOVA, followed by a Student’s t-test. In Figure S5, the significance of the differences between groups of samples was estimated with a Mann & Whitney non-parametric test, since the number of samples was less than 30 for each group. For correlation plots, the correlation coefficient was calculated with the cor function using the Pearson method and visualized using the corrplot function within the corrplot package, v0.84 (https://github.com/taiyun/corrplot). A heatmap was drawn with the plot_heatmap function of the ggplot2 package, v3.0.0 (Wickham, 2010).

#### Data availability

Raw paired sequences have been deposited in the Sequence Read Archive database under the accession numbers SAMN11405181 to SAMN11405379 and SAMN11936727 to SAMN11936732, corresponding to BioProject PRJNA532535.

## 3. RESULTS

### 3.1. Sampling strategy

Through a survey of a single production unit, we identified five processing steps as potential sources of bacterial contamination, with each step potentially able to induce variations in bacterial species diversity and population level. These steps were the transportation of raw meat from the slaughterhouse to the ham-producing factory, churning, drain-off after cooking, slicing, and packing. Although this is an industrial process, over the course of a year, these steps are often subject to important variations in duration or in exposure to aerobiosis/anaerobiosis conditions. To assess the role of each step in shaping bacterial diversity, we set up a complete factorial design in which each of the five processing steps was sampled at two different levels, chosen specifically from those encountered in the focal production unit (Table 1). With this sampling design, a total of 32 hams were sampled, each produced under slightly different conditions (Fig. 1). Although all meat was purchased from the same slaughterhouse, we also assessed possible variations due to changes in the initial meat microbiota over an extended period; specifically, we conducted four separate sampling campaigns over a period of six months: in June (Campaign C1), August (Campaign C2), September (Campaign C3), and November (Campaign C4). Overall, 128 different hams (32 × 4) were sampled.

**FIG 1.**
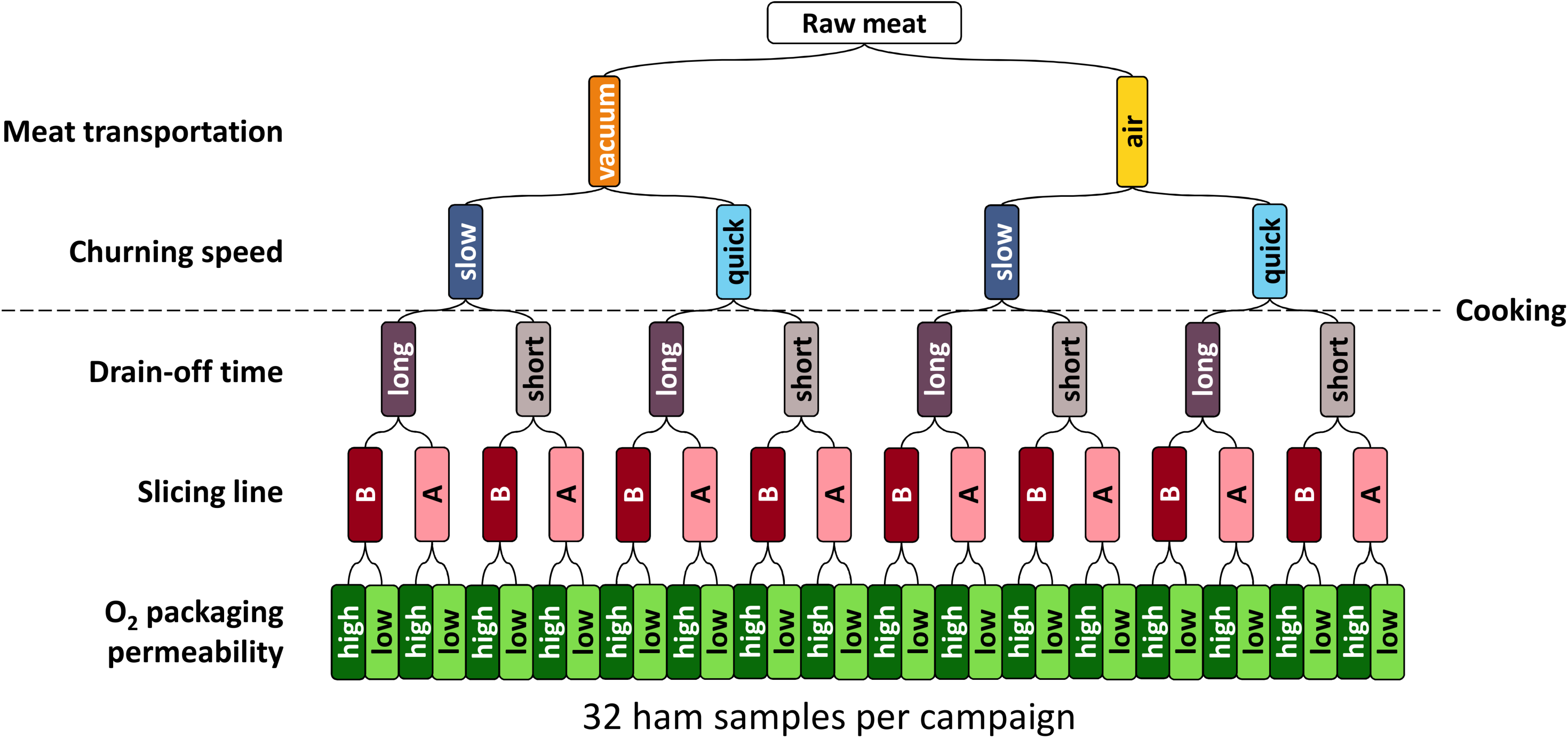
Ham sampling strategy. Sampling strategy based on a complete factorial design, with two levels for each of the five processing steps: meat transportation, churning, drain-off, slicing, and O_2_ packaging permeability.

The variations in ham production that were evaluated by our factorial design were necessarily accompanied by variations in the total length of processing, from a minimum of 12 days to a maximum of 28 days. However, optimal operational procedures for slicing and packing required that these two steps be carried out with the shortest delay possible. Due to these constraints, each sampling campaign required the use of four different batches of raw meat for ham production, as shown in Figure S1. Although our study was primarily focused on the final bacterial diversity of ham (after cooking, slicing, and packing), we incorporated some controls by measuring the bacterial diversity before cooking (after churning) in two campaigns (C2 and C3). Finally, each cooked ham was analyzed at two storage times: at the use-by date (UBD) and after 15 days of extended storage (past UBD), corresponding respectively to samples stored at 8°C for 28 and 43 days (see Fig. S1). Altogether, the sampling comprised a total of 256 hams.

### 3.2. Bacterial diversity in cooked ham was low and was shaped by both pre- and post-cooking processing steps

Bacterial diversity was assessed with MiSeq paired-end *gyrB* amplicon sequencing, which enabled accurate taxonomic assignment of OTUs to the species level (Poirier et al., 2018b). Amplification of 16S rRNA gene was performed for fourteen samples as a control and confirmed that there was no bias in species detection resulting from the use of *gyrB* (see supplementary Fig. S2). For *gyrB* amplicon sequencing, the success rate for DNA extraction and sequencing was around 80%, which yielded a dataset of 200 samples out of the 256 initially planned. However, the failed samples were scattered throughout the factorial design and campaigns and thus did not reduce the effectiveness of our statistical analysis. From the whole *gyrB* dataset, 157 OTUs were identified and assigned to 44 different bacterial species (average of four OTUs per species, see supplementary dataset S3 for a table detailing OTU abundance). As shown in Table 2, the mean number of species identified per sample was approximately 15 species.

**TABLE 2.** Comparative table showing the number of species (after merging OTUs with similar taxonomic assignments) identified in different subsets of samples

|  | Sample subsets |  |  |
| --- | --- | --- | --- |
|  | Churned meat<br>(15 samples) | Ham at UBD<br>(92 samples) | Hams past UBD<br>(108 samples) |
| **Campaign 1 \*** | - | 16 ± 3 | 13 ± 4 |
| **Campaign 2 \*** | 50 ± 8 | 13 ± 4 | 14 ± 5 |
| **Campaign 3 \*** | 41 ± 10 | 13 ± 6 | 14 ± 7 |
| **Campaign 4 \*** | - | 14 ± 7 | 25 ± 6 |
| **Total number of species** | 55 | 43 | 44 |
|  | 70 |  |  |

\* Mean number of species

The majority of the bacterial diversity in cooked ham samples was represented by approximately fourteen different species in various combinations, which together corresponded to a mean of 98% of the total relative bacterial abundance (Fig. 2). To correlate differences in bacterial community composition to variations in processing steps, we performed a comparative compositional analysis using the Bray-Curtis dissimilarity index and non-metric multi-dimensional scaling (NMDS). This analysis revealed that ham bacterial communities at the UBD were not significantly different from those past the UBD, and further revealed that slicing line and sampling campaign were the main factors that shaped these communities (Fig. 3). In order to better understand these differences, bacterial assemblages were plotted according to these two processing parameters. Figure 4 illustrates how these parameters influenced both the relative abundance and presence/absence of core bacterial species. Briefly, the identity of the slicing line influenced the overall distribution of multiple bacterial genera (e.g., *Carnobacterium* in line A *versus Leuconostoc* and *Serratia* in line B). Within each genus, the slicing line also influenced the identity of the species present (e.g., *C. divergens* in line A *versus C. maltaromaticum* in line B). Certain species also appeared in multiple sampling campaigns of a given line (e.g., *L. mesenteroides* present in campaign C1 and C2 for line A; *L. gelidum* subsp*. gelidum* present in campaign C3 for line B; *L. sakei* present in campaign C1 for line B; *C. maltaromaticum* present in campaign C2 and C3 for line B).

**FIG 2.**
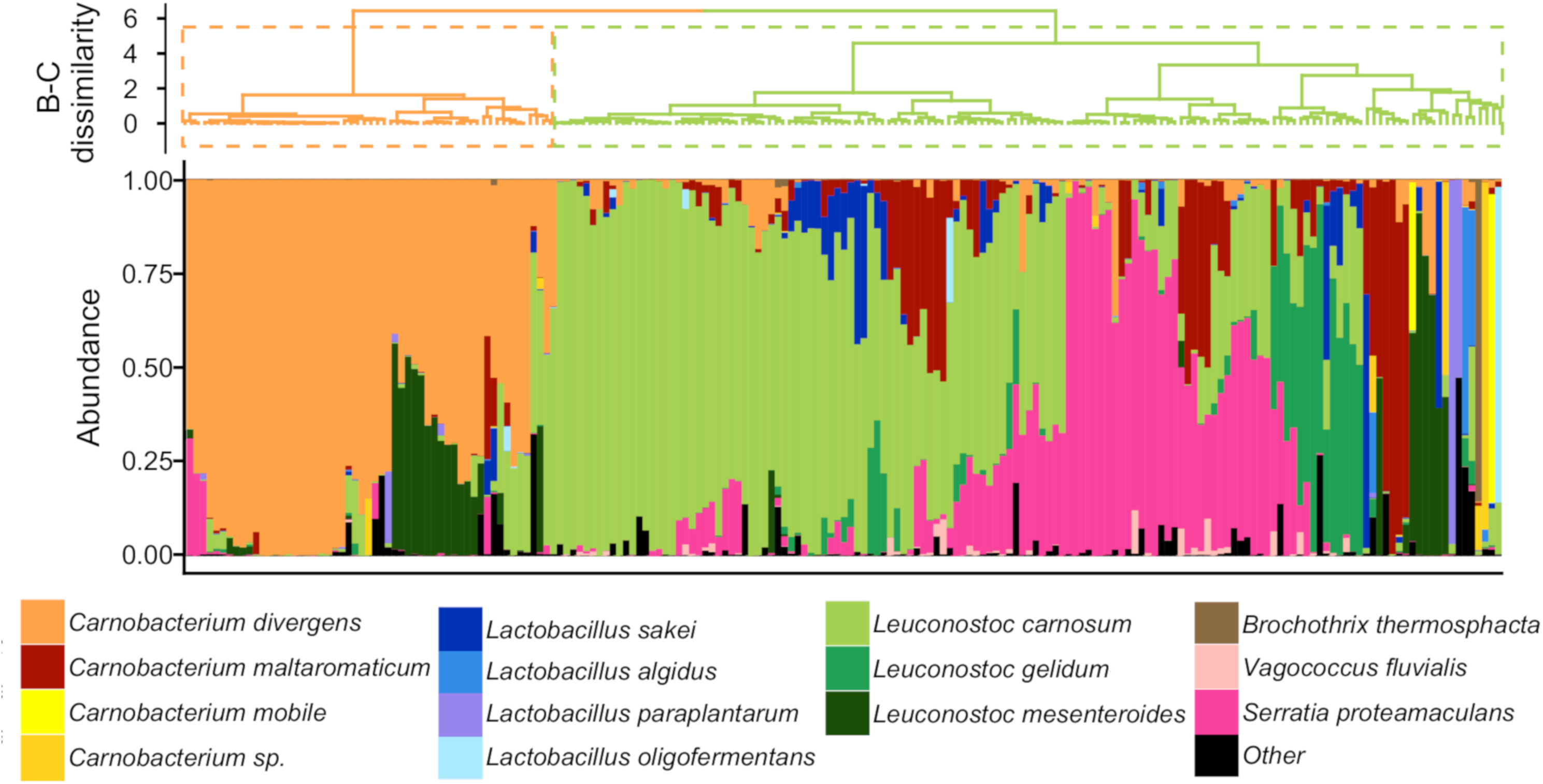
Ward unsupervised clustering of the microbiota of cooked ham samples according to Bray-Curtis dissimilarity index. The clustering analysis revealed two main clusters which were further subdivided into several clusters of various sample sizes. Below the clustering tree is a species composition plot (relative abundance) for the fourteen most-abundant species.

**FIG 3.**
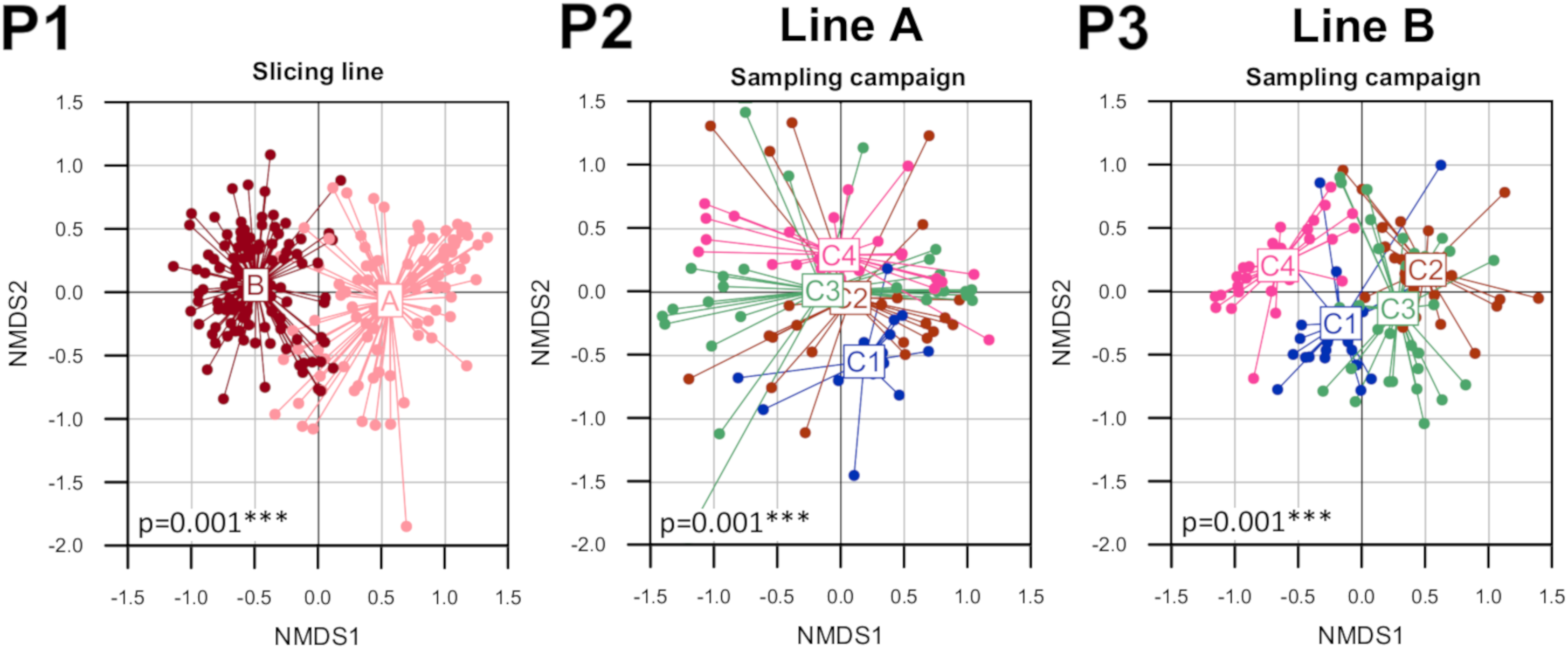
Non-metric multi-dimensional scaling analysis of cooked ham microbiotas based on Bray-Curtis dissimilarity index. Panel P1: The bacterial assemblage of each cooked ham sample is represented by a dot, colored according to the slicing line on which the ham was processed (red = slicing line B; pink = slicing line A). Panels P2 and P3: cooked ham assemblages were subdivided based on slicing line (most discriminant parameter), and each subset (line A in panel P2; line B in panel P3) was analyzed separately, with the samples colored according to their sampling campaign of origin (blue = campaign C1; red = campaign C2; green = campaign C3; pink = campaign C4). The other processing steps had no influence in structuring overall bacterial diversity (see supplementary Fig. S4). For slicing line and sampling campaign, differences in composition between the bacterial communities present in the tested conditions were estimated using permutational ANOVA. Corresponding p-values are indicated under each plot.

**FIG 4.**
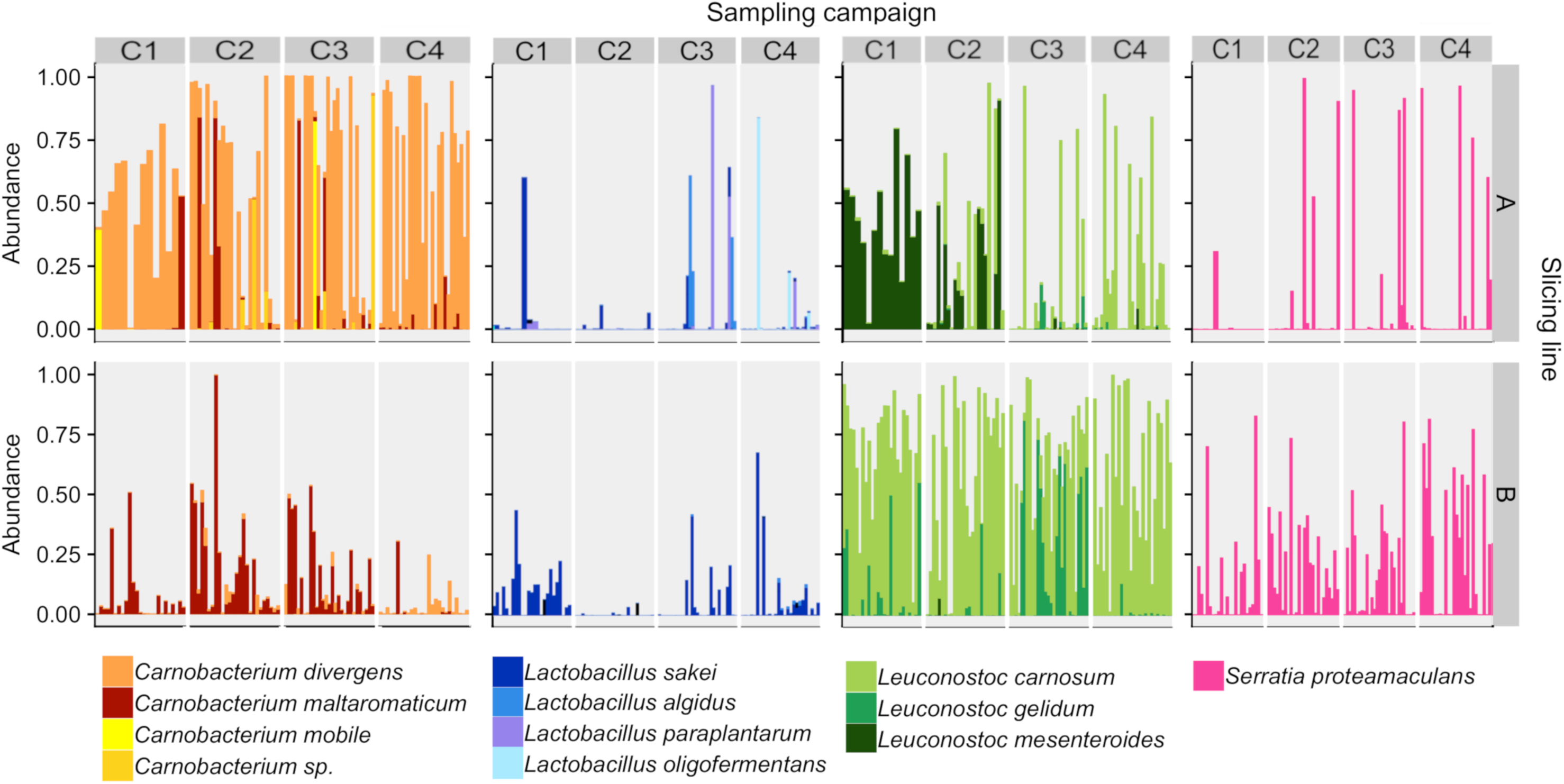
Species composition plots of cooked ham microbiota separated by slicing line and sampling campaign. Bar plots show the relative abundance of the fourteen core bacterial species in cooked ham. Cooked ham samples are distributed vertically by slicing line of origin (top = samples from line A; bottom = samples from line B). Within each slicing line, ham samples are distributed horizontally according to the four main bacterial genera present (from left to right: *Carnobacterium, Lactobacillus, Leuconostoc*, and *Serratia*) and then according to sampling campaign (C1 to C4).

It should be noted that the effect of churning time on the cooked ham microbiota was also revealed to be statistically significant, although it was less obvious from the NMDS analysis (see supplementary Fig. S4). Indeed, this effect was mostly significant in cooked ham from slicing line A. Similarly, O_2_ packaging permeability had a significant effect on the microbiota of line B only. We linked this observation to the fact that *S. proteamaculans,* the only dominant Proteobacteria found among cooked ham samples, was much more prevalent in samples from slicing line B (in 58% of samples from line B *versus* 17% of samples from line A). Moreover, samples of line B differed greatly in the relative abundance of this species, which suggests the influence of another processing step than the slicing line. Indeed, when samples from line B were analyzed separately, we observed that *S. proteamaculans* was significantly more abundant in samples produced with processing steps that had higher oxygen levels (air transportation and high O_2_ packaging permeability; see Fig. 5 Panel P1). We also noted that, at the UBD, O_2_ was not detected in cooked ham samples packed in highly permeable packaging (data not shown). Therefore, any O_2_ that permeated the packaging seemed to have been rapidly consumed by the microbiota. In slicing line A, instead, the relative abundance of *C. divergens* appeared to be correlated with churning, as the abundance of this species increased in ham that was churned quickly (see Fig. 5 Panel P2).

**FIG 5.**
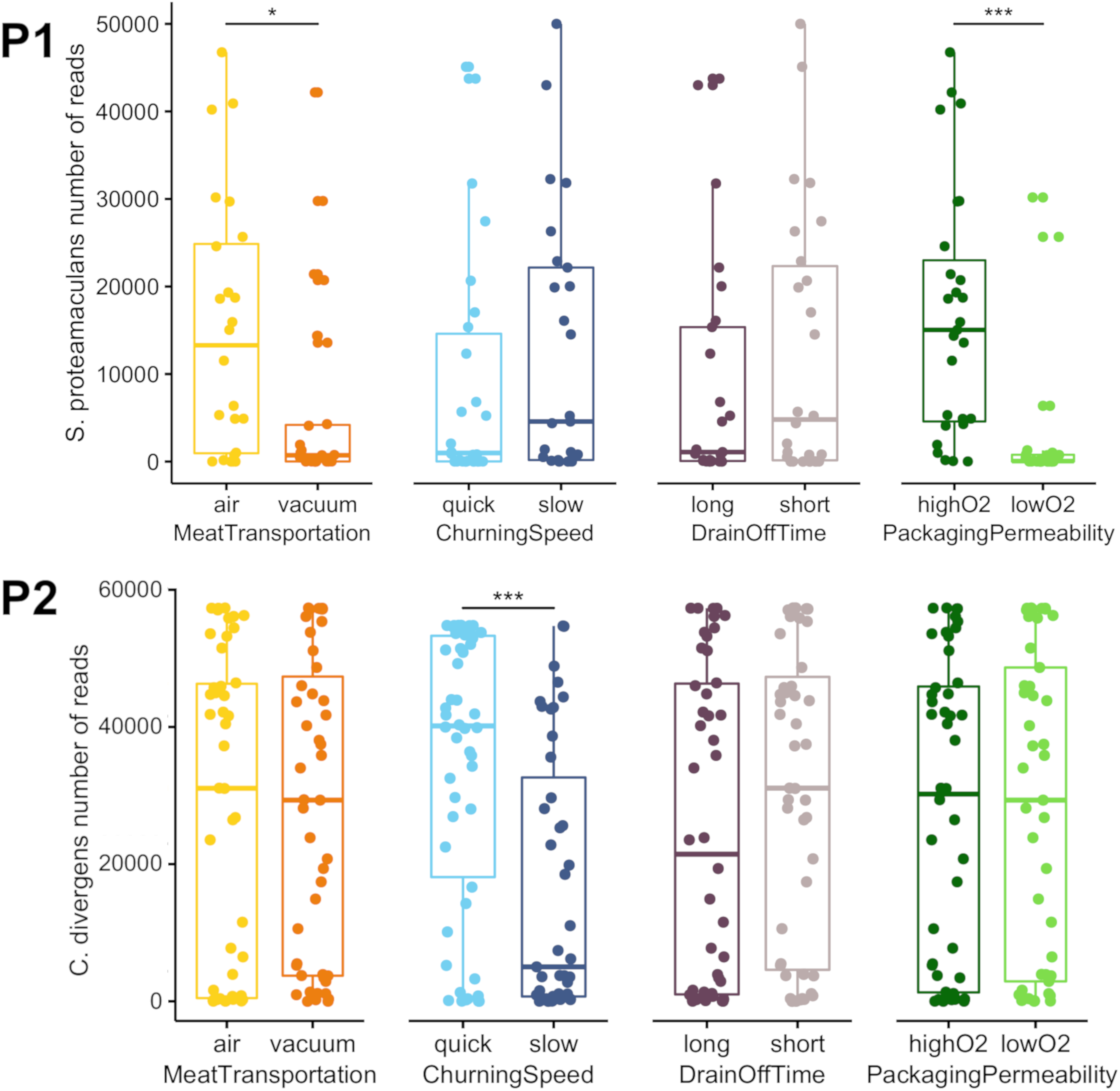
Comparative analysis of the relative abundance of *S. proteamaculans* and *C. divergens* in cooked ham samples as a function of processing parameters. Panel P1: relative abundance (in number of reads) of *S. proteamaculans* in samples from slicing line B as a function of the other processing parameters. Panel P2: relative abundance (in number of reads) of *C. divergens* in samples from slicing line A as a function of the other processing parameters. Statistical variance between groups of samples was determined with ANOVA followed by a Student’s t-test. The number of stars indicate the p-value: *** p-value < 0.001; ** p-value < 0.01; * p-value < 0.05.

To summarize, our data indicated that the four main processing parameters that shaped the types of bacterial communities present in cooked ham were, in decreasing order of influence, identity of slicing line, sampling campaign (different meat batches over six months of sampling), O_2_ permeability of packaging, and churning speed. Two of these processing steps occurred post-cooking (slicing line and packaging type) and two occurred pre-cooking (campaign and churning speed).

### 3.3. Bacterial species present in cooked ham mostly originated from raw meat and were among the most abundant species in meat samples after churning

Because the slicing line had a strong influence on the type of microbiota present in ham samples, even in samples that originated from the same batch of raw meat, we next investigated whether the bacterial species identified after cooking were also present before cooking. Furthermore, the slight influence of pre-cooking factors such as sampling campaign and churning speed on the relative abundances of certain species raised questions regarding the effectiveness of the cooking step in controlling bacterial populations. As shown in Fig. 6, we found that the most abundant species in cooked hams were also present in raw meat after churning: *C. divergens, L. sakei*, *C. maltaromaticum*, *L. carnosum, L. gelidum*, *S. proteamaculans*, and *L. mesenteroides*, in decreasing order of abundance. Overall, cooking eliminated only 50% of the species present in raw churned meat, mostly those that were subdominant. Among species that were highly abundant in raw churned meat, the cooking step eliminated Proteobacteria (e.g., *Pseudomonas* species) more effectively than Firmicutes, with the exception of *S. proteamaculans.* However, we also found evidence for post-cooking contamination, as illustrated by the presence of species in cooked hams that had not been detected in raw churned meat for those sampling campaigns (e.g*., Vagoccocus fluvialis*).

**FIG 6.**
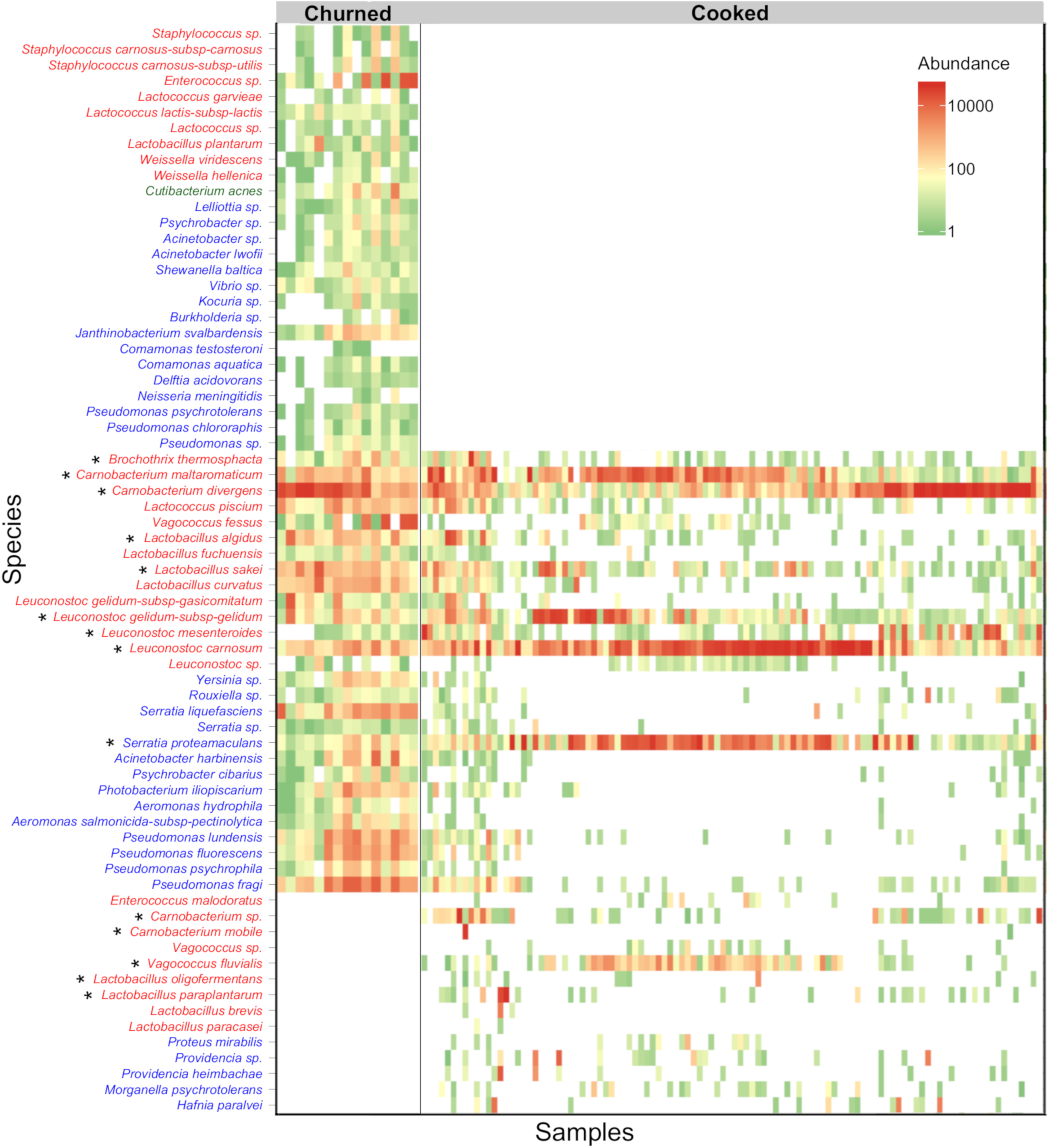
Heatmap depicting the effect of cooking on bacterial species abundances in ham. Samples from campaigns C2 and C3 are ordered according to the sample type: left, raw meat after churning, and right, cooked ham at UBD. Bacterial species are ordered in three groups from top to bottom: species present only before cooking (hence eliminated by the cooking step), species present before and after cooking (either resistant to cooking or reintroduced after cooking), and species present only after cooking (contamination). Within each group, species are ordered by their taxonomic relationships. Species names colored in red, blue, or green indicate placement within the phyla Firmicutes, Proteobacteria, or Actinobacteria, respectively. Stars indicate the fourteen most-abundant species in cooked ham. The scale in the top-right corner of the heatmap depicts the color palette associated with the relative numbers of reads of the various species. The black vertical line represents the cooking step.

### 3.4. The production of fermentation-derived metabolites, used as spoilage indicators, was species-dependent and driven by population abundance

Next, we focused our analysis on the consequences of these different community patterns for the quality of the cooked ham. Of the important characteristics of spoilage, we chose to assess bacterial load, decrease in pH (an indicator currently used in the meat industry to detect spoilage), and the production of volatile organic compounds (VOCs). Additionally, since ham contains glucose, we also measured the concentration of end-products from the mixed-acid fermentation of pyruvate (both L- and D-lactic acid, acetic acid, formic acid, and ethanol). Because no differences were detected in the bacterial communities at the two sampling times (UBD or past UBD), we narrowed our analysis to UBD samples only (*n=*128) in order to determine whether any significant differences in spoilage characteristics were detectable in hams at the end of storage, prior to consumption.

At the UBD, the bacterial load of cooked hams from slicing line A (7.1 ± 1.0 log_10_ cfu.g^−1^) was not significantly different from that of hams from slicing line B (7.7 ± 0.6 log_10_ cfu.g^−1^). However, there were notable differences among ham samples, with a range of 4 to 9 log. Furthermore, we were intrigued that at a similar bacterial load, the mean decrease in pH from immediately after packaging (6.15 ± 0.06) to the UBD (6.05 ± 0.15) was significantly larger in samples from slicing line B (Fig. S5) and only when the bacterial load reached 7 log_10_ cfu.g^−1^. This prompted us to analyze the possible correlation between decrease in pH, production of volatile organic compounds, pyruvate-derived end-products, and population abundances of the different bacterial species present in the microbiota of the two slicing lines.

From an initial screening of volatile organic compound production in hams, we noticed that the majority of VOCs were pyruvate-derived end-products, with ethanol representing around 90% of the total volatilome (see supplementary data Fig. S6), followed by butan-2,3-dione and butan-2-one in smaller proportions. Compounds derived from amino-acid metabolism, such 2-methylbutanal and 2-methyl-1-butanol, were also detected, but represented less than 5% of the total volatilome. Because of this, we focused our subsequent analyses on the main pyruvate-derived end-products.

As shown in Fig. 7, an analysis of Pearson correlations revealed that decrease in pH was positively correlated to the production of D-lactic acid, acetic acid, and formic acid. The production of D-lactic acid was strongly correlated to that of ethanol and to the population abundance of *Leuconostoc* species (mainly *L. carnosum*). This observation is also illustrated in supplementary data Figure S7 showing the relationship between the production of end-products and the population abundance of each of the bacterial species that were targeted as potential producers. Instead, the production of acetic acid also showed a strong positive correlation with that of formic acid and to many of the species that are co-occurring on ham from slicing line B; namely *S. proteamaculans, C. maltaromaticum* and *V. fluvialis* (and to a lesser extent with *B. thermosphacta*). *C. maltaromaticum* in particular showed the strongest correlation to the production of formic acid. As presented earlier in Fig. 4, *L. carnosum*, *C. maltaromaticum*, and *S. proteamaculans* were the most dominant species in samples from slicing line B, indicating that this type of microbiota may have a higher potential for spoilage than that of slicing line A. Conversely, the bacterial community of ham samples from slicing line A, in which these three species were subdominant with respect to *C. divergens,* correlated to lower production of these compounds. The production of L-lactic acid, on the other hand, was correlated to the abundance of *L. sakei* and *C. mobile*.

**FIG 7.**
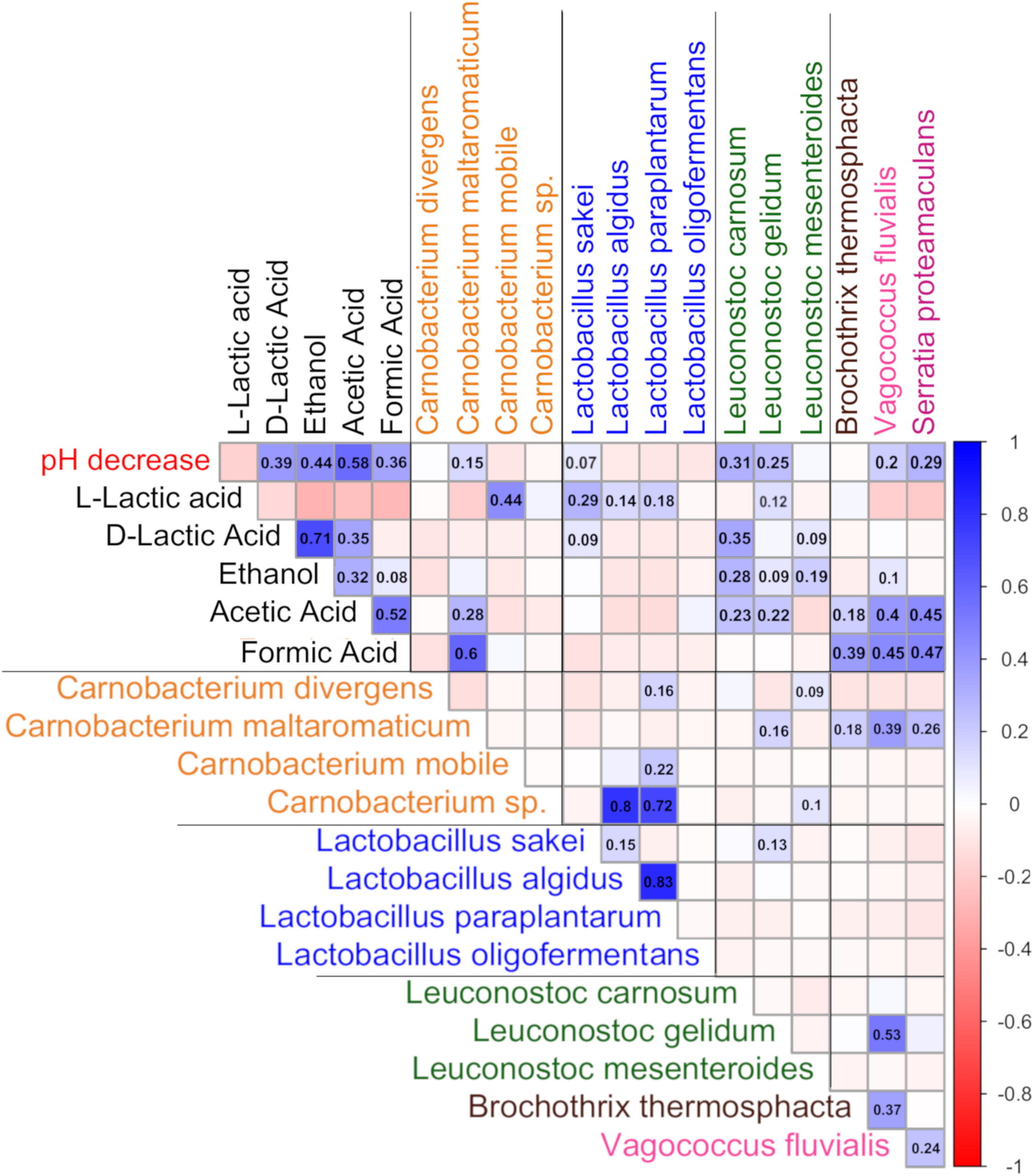
Correlation plot representing the Pearson coefficient between the production of end-products from the mixed-acid fermentation of pyruvate and the estimated load of bacterial species on cooked ham at the UBD. Fermentation end-products are vertically distributed according to the difference between the final concentration at the UBD and the initial concentration just after packaging. The absolute load of each bacterial species is distributed horizontally and was estimated using the total bacterial load weighted by each species’ relative abundance (percentage; see Materials & Methods). The Pearson correlation coefficient between the production of an end-product and the estimated load of a bacterial species is represented by a colored square. The scale on the right depicts the color palette associated with the value of the Pearson correlation coefficient (red, white, and blue corresponding respectively to a negative correlation, no correlation, and a positive correlation).

These results also prompted us to verify the effect of post-cooking factors (slicing line and O_2_ packaging permeability) on the metabolic indicators (see Fig. 8). With the exception of L-lactic acid, all compounds were present in higher concentrations in cooked ham from slicing line B. Furthermore, changes in the O_2_ permeability of the packaging resulted in a switch from L-lactic acid production (low O_2_) to acetic acid production (high O_2_).

**FIG 8.**
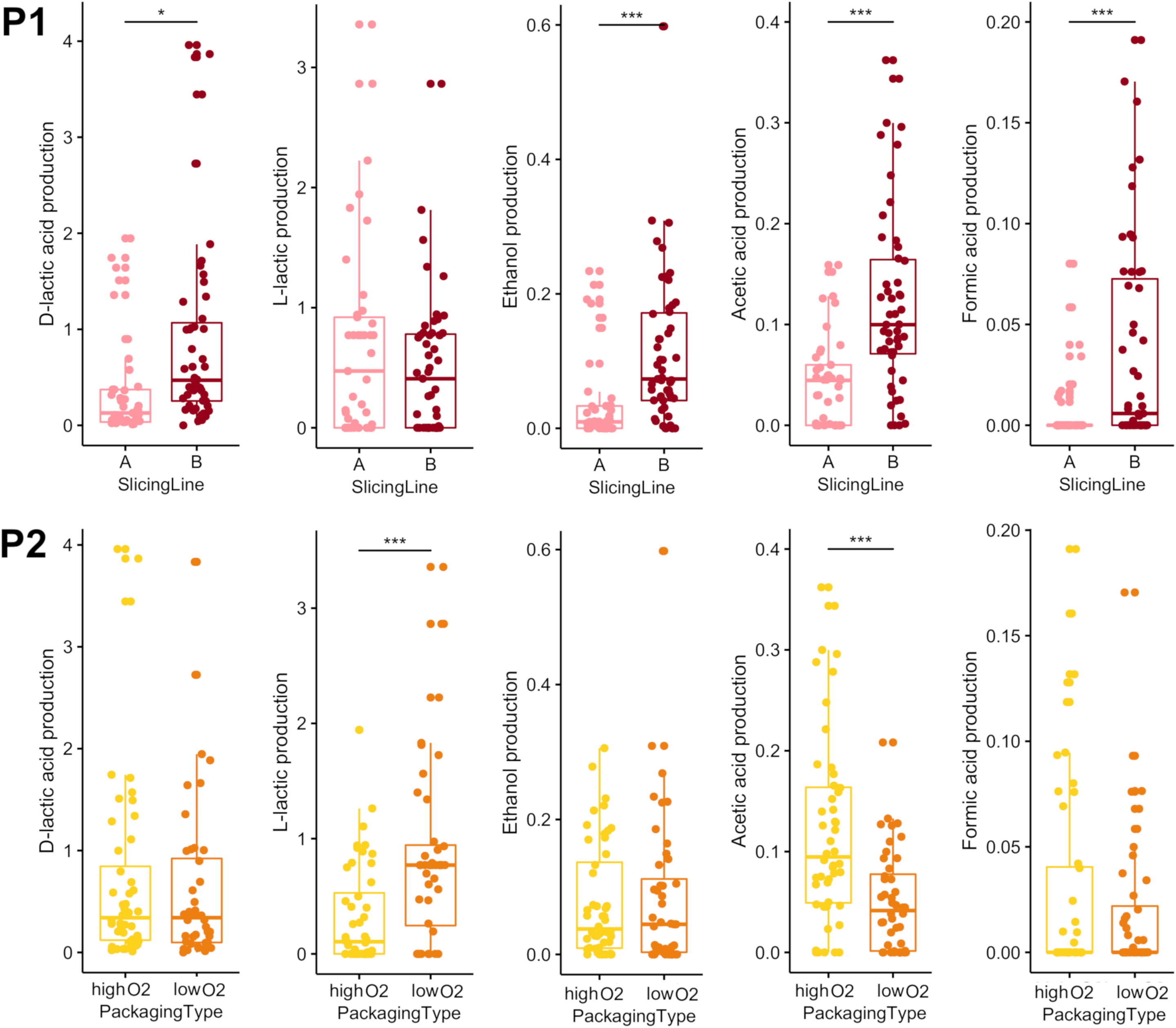
Box plots of the production in mg.g^−1^ of end-products from the mixed-acid fermentation of pyruvate as a function of the estimated load of bacterial species at the UBD. For each graph, all UBD samples were sorted according to their estimated load of a species of interest within the bacterial community (e.g., samples were sorted according to their estimated load of *C. maltaromaticum*). Samples for which the estimated load of the species of interest was less than 2 log_10_ cfu.g^−1^ were not included in order to focus on samples in which the species was relevant. In some cases, high bacterial loads were not attributable to an identified species and were labeled “NA”. *B. thermosphacta* and *C. mobile* were not included because they were present in too few samples to allow box plot representation. Panels depict the relationships for different metabolites, from P1 (D-lactic acid) to P5 (formic acid), presented by decreasing average production levels.

## 4. DISCUSSION

The aim of this study was to perform a large-scale analysis of bacterial diversity in order to survey and characterize the influence of the processing steps in the microbial ecology of cooked ham. Among the various parameters described in previous studies and known to play a role in the variability of this food product, we have chosen five of them upstream or downstream of the cooking step. To do this, we focused on a single facility and deployed an extensive sampling strategy, with a complete factorial design and we applied it several times over four months of production to strengthen our findings. Our study shows evidences for the role of major and minor factors on both sides of the cooking steps that are driving the microbial ecology of cooked ham and hence the ham quality.

### 4.1. After cooking, the slicing step creates most of the variability in ham bacterial communities and shapes the development of different spoilage metabolic profiles

Our first unambiguous demonstration is that, the slicing line was the main source of variability in the microbial assemblages of ham. Although similar implication of the slicing equipment in bacterial contamination has been previously made by other scientists (Pothakos et al., 2012; Vasilopoulos et al., 2008), our analysis provides a better view of the impact that this process parameter is playing in ham quality. Indeed, in addition to prove that ham samples originating from the same batch of raw churned meat ended up with different bacterial communities based on the slicing line on which they had been cut after cooking, we demonstrate that each slicing line has a proper microbial fingerprint which remains rather stable in time. In the present case, the time spanned over several months from June to November during which the slicing lines were used several hundreds of times in the production unit between each sampling campaign. Furthermore, we show that the quality of cooked hams is also impacted by this specific processing step. It is important to remember here that the cooked ham samples that we analyzed at the UBD were generally not spoiled, and the production of spoilage metabolic indicators was usually very low. However, samples from line B showed higher production levels of fermentation-derived spoilage metabolites and a stronger pH decrease than those from line A. Unsurprisingly, production of D-lactic acid and ethanol was positively correlated with the prevalence and population abundance of species from the genus *Leuconostoc* (especially *L. carnosum*) and thus was more elevated in samples from line B. Such finding corroborates previous studies which showed that members of the *Leuconostocaceae* family are generally identified as strong putative spoilers, since their presence in high amounts in meat samples is often associated with gas production, greening, ethanol production and/or acidification (Björkroth et al., 1998; Comi and Iacumin, 2012; Samelis et al., 1998; Raimondi et al., 2019). This finding is also consistent with two aspects of *Leuconostoc* biology. Firstly, the well-known heterolactic fermentation of glucose by species belonging to the *Leuconostocaceae* family under anaerobiosis (Cogan and Jordan, 1994; Dols et al., 1997) leading to lactate, acetate and ethanol as main end products. Secondly, the fact that *Leuconostoc* species preferentially produce D-lactate due to several D-lactate dehydrogenase-encoding genes in their genome.

Our results allowed us to draw similar conclusions on the impact of the slicing line on the production of acetic acid and formic acid (both with high correlation to pH decrease), increased in hams from line B when compared to hams from line A. The production of acetic acid by the ham microbiota is more challenging to interpret because, in addition to *Leuconostocs* species, several other species found on samples from line B can be involved in this process, most of which demonstrate co-occurrence relationships, such as *C. maltaromaticum*, *B. thermosphacta, S. proteamaculans*, and *V. fluvialis*. Nevertheless, our data are pointing out that the slicing line, as the main influencing factor on the microbiota structure, is modifying the production profile of these two acids. Thus, our results show that *C. maltaromaticum* and *S. proteamaculans* are the second confounding spoilers (after *Leuconostocs*) of cooked ham as they correlate with accumulation of acetic and formic acids. In particular, these two species distinguish themselves by specific production of formic acid in case of *C. maltaromaticum*, corroborating previous observations on *Carnobacterium* metabolism in meat (Borch and Molin, 1989; Zhang et al., 2019); and acetic acid by *S. proteamaculans.*.

One of the reasons we suspect this *Serratia* species of being involved in acetic acid production is the influence played by the packaging with high O_2_ permeability on both parameters, thereby indicating that this processing factor is also playing a role in the microbial ecology of cooked ham. This post-cooking condition favored both higher concentrations of acetic acid and the growth of *S. proteamaculans* in samples sliced on line B, while it did not affect microbial communities of hams sliced on line A (mostly deprived of *S. proteamaculans*). It is known that this facultative anaerobic species grows preferentially in the presence of O_2_, as shown in a previous study of batch culture that compared growth with and without O_2_ (Alfaro et al., 2013). However, we were not able to detect oxygen in the packages of cooked ham at the UBD, indicating that when it permeates through the packaging, it is rapidly consumed by the microbiota and that *S. proteamaculans* alternates between quick aerobic respiration (when O_2_ permeates the packaging) and longer period of glucose fermentation to acetate (when the O_2_ is depleted).

### 4.2. Raw meat microbiota is another source of bacterial variability between samples and has a sampling campaign effect

Our data revealed that the composition of the cooked ham microbiota also depended on pre-cooking parameters, in particular on the sampling campaign (different meat batches), but this effect was somewhat dependent of the slicing line and only specific groups of species are involved in this variability such as (*e.g. L. mesenteroides* and *L. sakei* in June or *L. gelidum* in September). Seasonal variability of raw meat microbiota has already been evidenced (Lucquin et al., 2012). However, our observation that a campaign-based effect can be evidenced on a cooked product raises the issue of the cooking step’s effectiveness in eliminating initial meat microbiota. Indeed, such observation has already been done on cooked pork sausages (Hultman et al., 2015). Here, we clearly show that the cooking step eliminated about half of the bacterial species present, mainly those with low abundance on raw churned meat, such as *Staphylococcus*, *Pseudomonas,* and *Yersinia*; whereas some species were clearly introduced during production (re-contamination after cooking), such as *V. fluvialis*. Most interestingly, we show that the dominant bacterial species identified on cooked ham at UBD could be traced before and after cooking. We interpret this result as being the likely consequence of two phenomena : (1) there is a persistence of post-cooking living cells which could explain, for instance, why air raw meat transportation would influence the prevalence and abundance of *S. proteamaculans* or why churning speed would influence the relative abundance of *C. divergens* on cooked ham at UBD; (2) the slicing line is either a source of selection or a further source of contamination. Both assumptions cannot be refuted at this time, as the microbial ecology of the slicing lines was not included in this study. The two slicing lines studied in the factory are equipment from two different generations. Surprisingly, the newer line B of which the cutting mechanism is more complex than that of line A, produces ham with higher spoilage potential.

## 5. CONCLUSIONS

Our extensive survey of the microbiota of cooked ham from a single production unit provided us with a detailed view of the microbiota variability over a long production time and to characterize the ways in which the microbial ecology of this product is shaped by the various processing steps. Our results clearly demonstrate that the process (and especially the slicing step) has a more important microbial fingerprint than the raw meat on the final communities of cooked ham. This study illustrates the very important role this type of strategy could play in the food industry if applied more broadly to process management, such as for instance, defining more precise UBD according to the conditions of production.

## LEGENDS TO THE FIGURES

**FIG S1 Management of sampling to optimize operational process requirements.**

The ham production schedule for a single campaign is shown as a function of variations in the length of the different steps (depicted in days at the top of the figure). After slicing and packing, ham samples (represented by red and yellow ellipses) were collected for all campaigns (C1 to C4), whereas meat samples were collected before cooking (represented by green ellipses) for campaigns C2 and C3 only.

**FIG S2 Principal Coordinates analysis based on Bray-Curtis dissimilarity index to compare ham microbial communities of samples using *gyrB* and 16S rDNA amplicon sequencing.**

Samples S1 to S14 are labeled “gyrB” or “16S” according to the molecular marker used for bacterial identification. Samples from the four different sampling campaigns (C1 to C4) are represented: S1 from campaign C1; S2 to S4 from campaign C2; S5 to S10 from campaign C3; S11 to S14 from campaign C4. Samples S1, S4, S5, S8, S10 and S13 were sliced on line A and samples S2, S3, S6, S7, S9, S11, S12 and S14 on line B.

**FIG S4 Non-metric multi-dimensional analysis of cooked ham samples based on Bray-Curtis dissimilarity index.**

Samples were analyzed in two separated subsets, corresponding to samples from line A on the left side and samples from line B on the right side. Within a given subset of samples, ham samples were colored either according to: the sampling campaign (blue=C1; red=C2; green=C3; pink=C4); the churning speed (dark blue=slow; light blue=quick); the O_2_ packaging permeability (dark green=high O_2_; light green=low O_2_); the conditions of transportation of raw meat (yellow=air; orange=vacuum); the drain-off time after cooking (dark grey=long; light grey=short); the shelf-life (light purple=UBD; dark purple=past UBD). For each graph, differences in composition between bacterial communities present in the two tested conditions were estimated using permutational ANOVA. Corresponding p-values are indicated under each plot.

**FIG S5 Boxplots depicting the decrease in pH of ham samples from slicing lines A and B as a function of total bacterial load.**

Bacterial load is represented in log_10_ cfu.g^−1^. Samples were segregated by their slicing line of origin and then sorted by their bacterial load, ranging from 4 to 9 Log_10_ cfu.g^−1^. Delta pH depicts the difference between the initial pH measured just after packaging and the pH at the UBD. Statistical variance between groups of samples within a slicing line was determined with a Mann & Whitney test. The number of stars indicates the p-value: * p-value < 0.05; “ns” for p-value > 0.05.

**FIG S6 Boxplots showing the percentage of total emitted volatile compounds of hams at UBD.**

Samples from campaigns C1 and C2 are represented. The compounds are ordered from left to right in decreasing order of abundance.

**FIG S7 Boxplots showing the production level in mg.g^−1^ of end products from mixed-acid fermentation of pyruvate according to the estimated load of bacterial species at UBD.**

For each graph, all UBD samples were sorted according to their estimated load of a species of interest within bacterial communities (e.g. samples are sorted according to their estimated load of *C. maltaromaticum*). Samples where the estimated load of the species of interest was under 2 log_10_ cfu.g^−1^ were not represented, in order to focus on samples where the species was representative. For some species, high loads were non-attributed and were labeled “NA”. *B. thermosphacta* and *C. mobile* were not represented since they were present in too few samples to allow boxplot representation. Metabolites are ordered in Panels, from P1 (D-lactic acid) to P5 (formic acid) by average decreasing production levels.

## Supporting information

Supplementary figures for Zagdoun et al, 2019

## ACKNOWLEDGMENTS

We thank the MIGALE bioinformatics platform at INRA (http://migale.jouy.inra.fr) for providing computational resources and data storage. We also thank the INRA @BRIDGe platform for carrying out the MiSeq sequencing runs. Marine ZAGDOUN was the recipient of ANRT grant number 2016/1038 (Association Nationale de la Recherche et de la Technologie, Ile de France). The funders had no role in the study’s design, data collection, and interpretation, or the decision to submit the work for publication.

