## Supplementary figures and images for "LARGE MICROBIOTA SURVEY REVEALS HOW THE MICROBIAL ECOLOGY OF COOKED HAM IS SHAPED BY DIFFERENT PROCESSING STEPS"

### Supplementary figures for Zagdoun et al, 2019

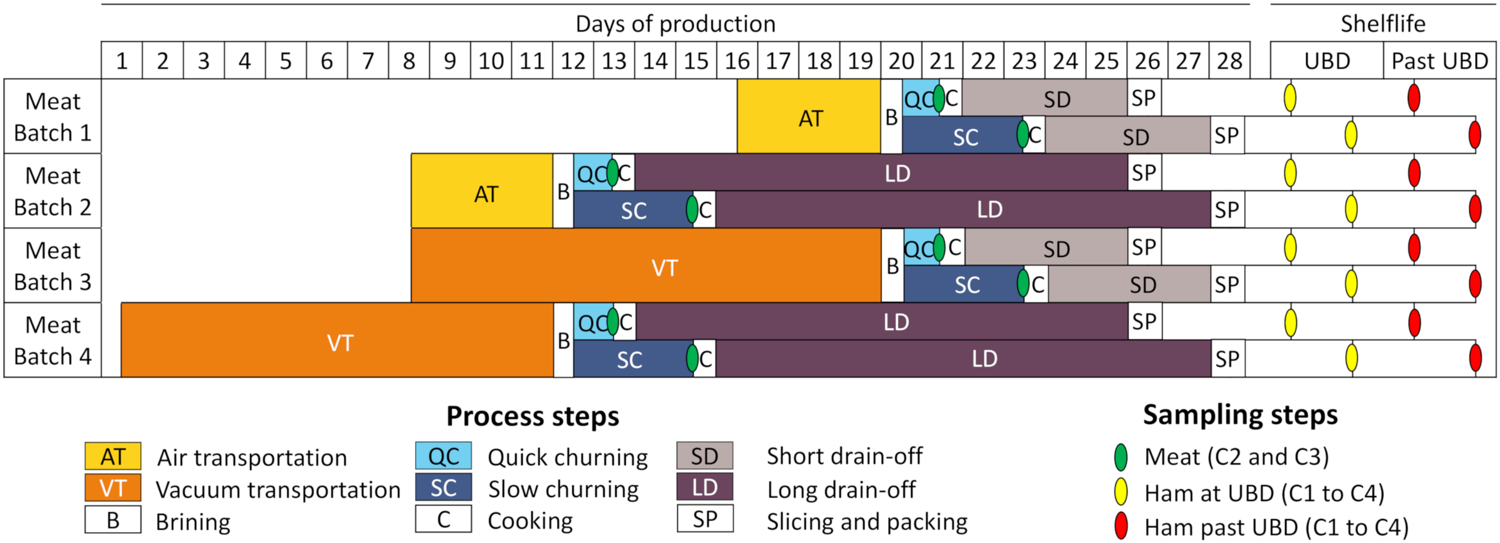

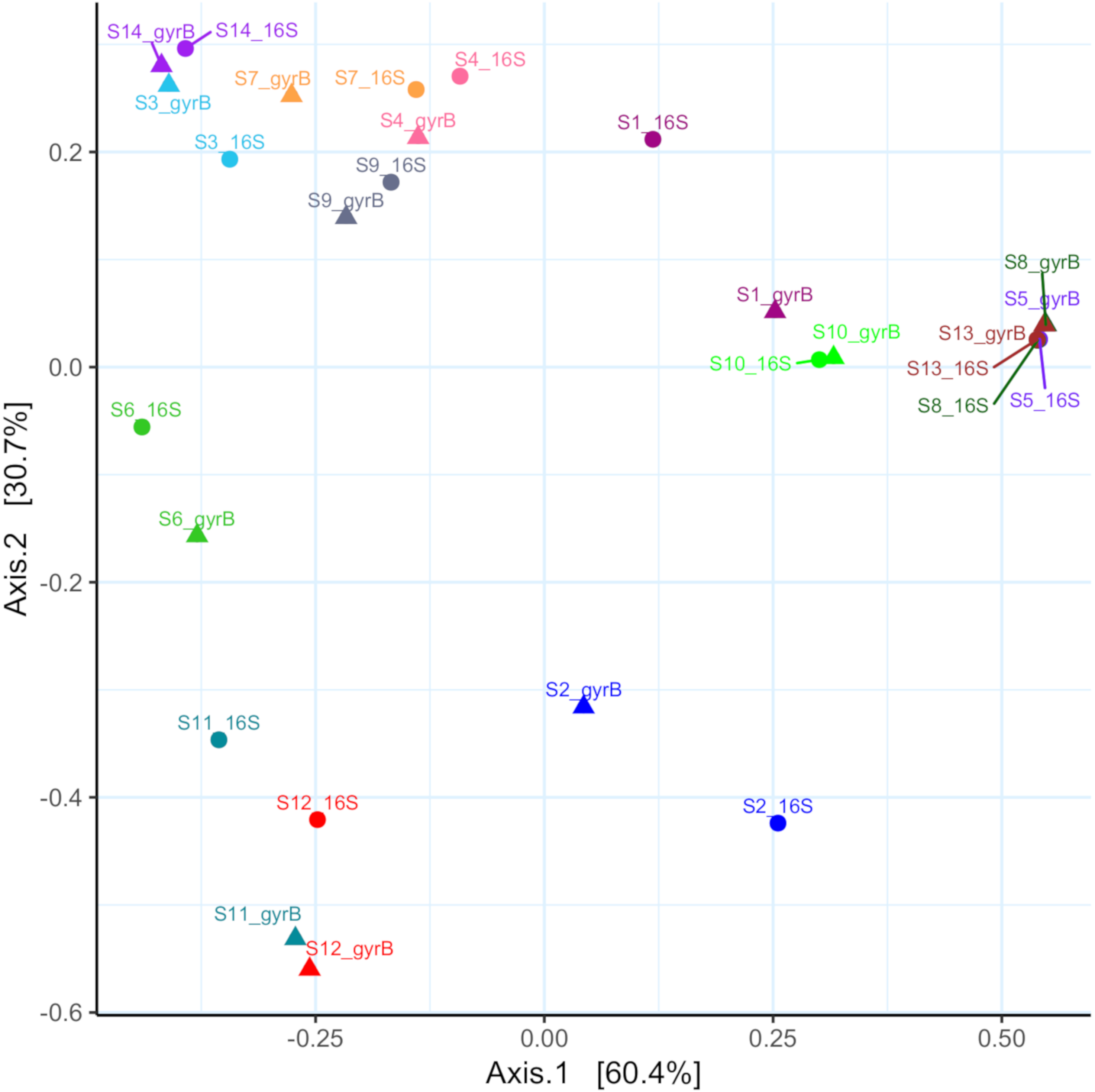

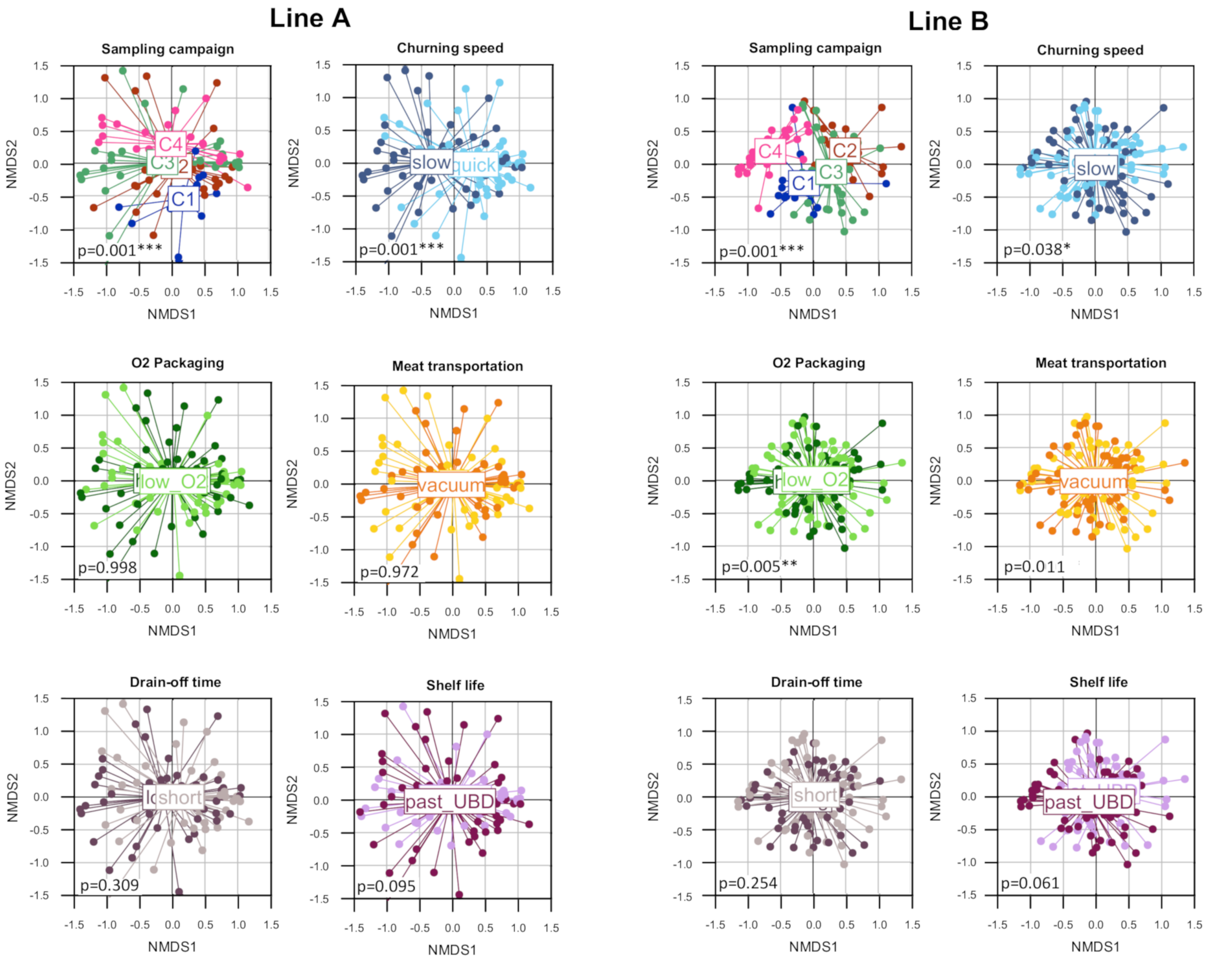

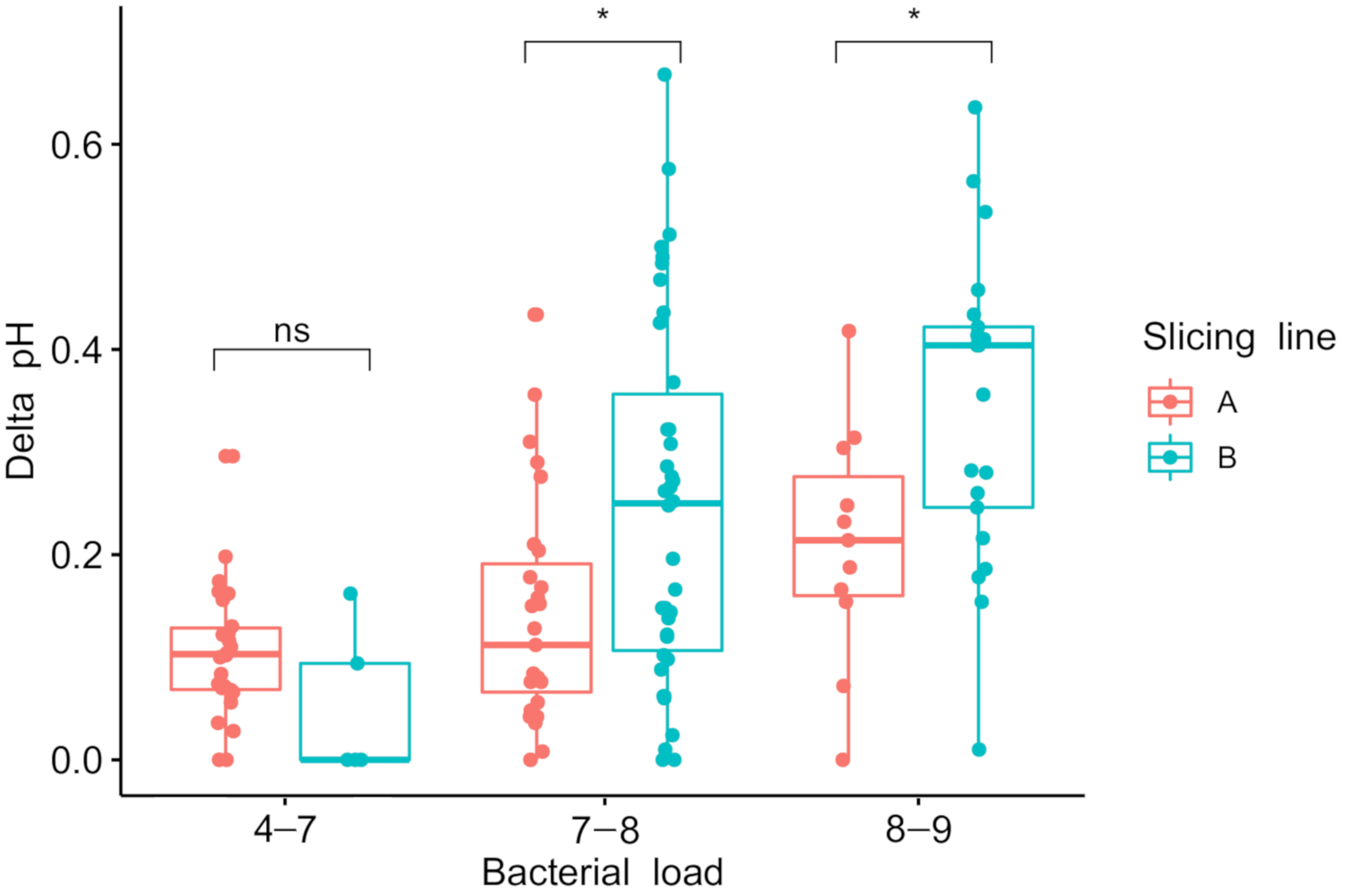

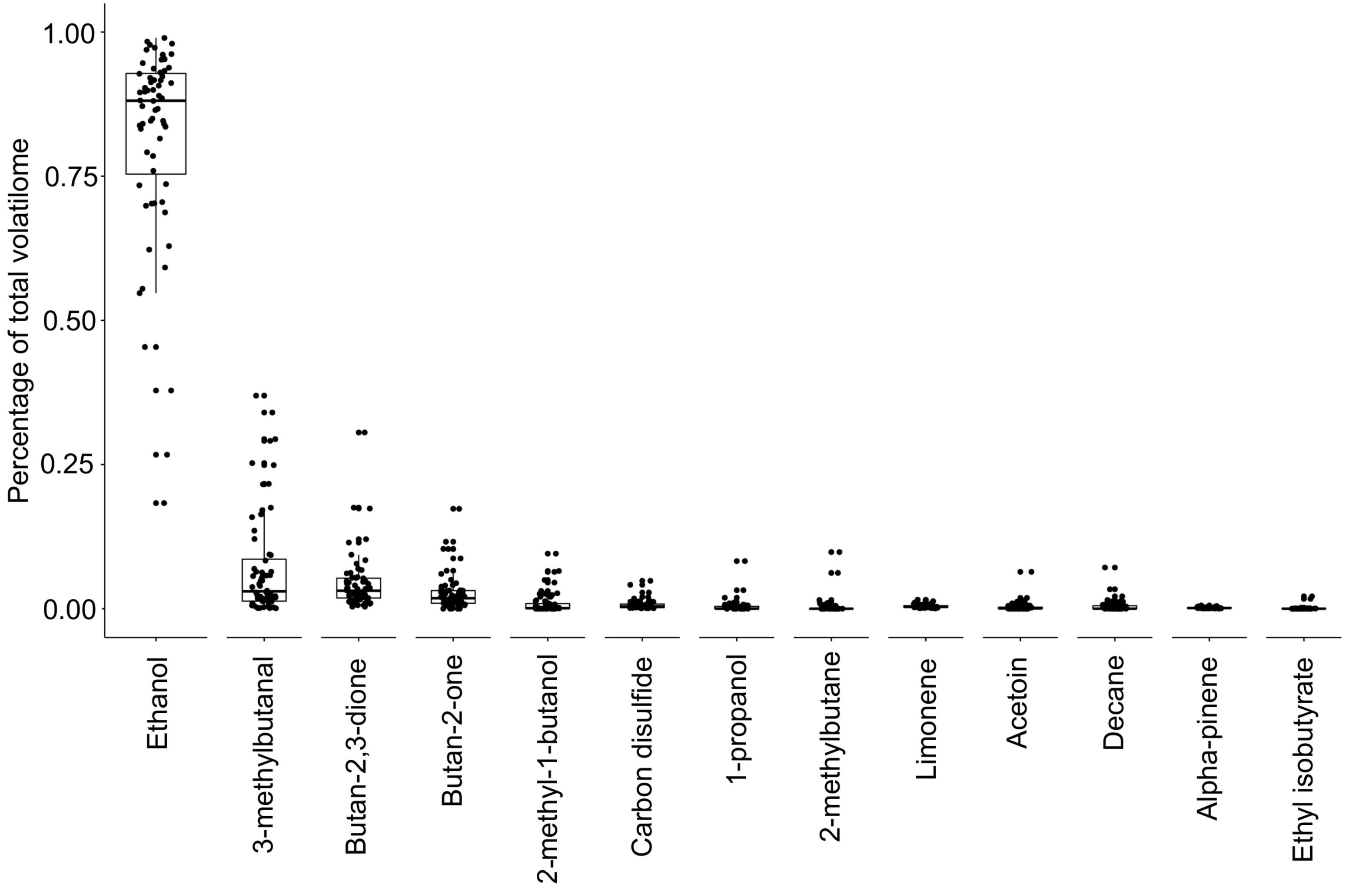

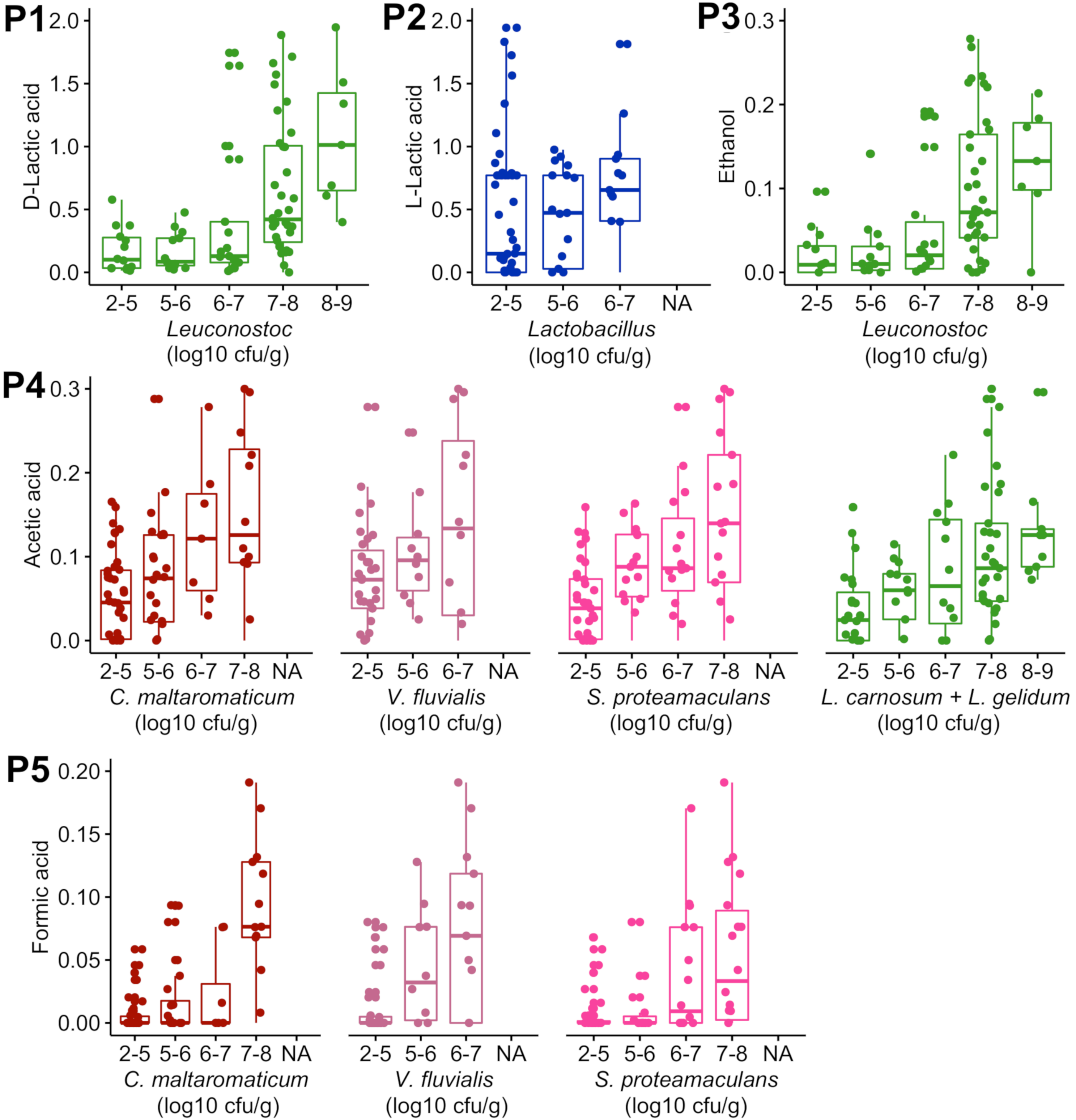
